# Attentional Modulation of the Auditory Steady-State Response across the Cortex

**DOI:** 10.1101/836031

**Authors:** Cassia Low Manting, Lau M. Andersen, Balazs Gulyas, Fredrik Ullén, Daniel Lundqvist

## Abstract

Selective auditory attention allows us to focus on relevant sounds within noisy or complex auditory environments, and is essential for the processing of speech and music. The *auditory steady-state response (ASSR)* has been proposed as a neural measure for tracking selective auditory attention, even within continuous and complex soundscapes. However, the current literature is inconsistent on how the ASSR is influenced by selective attention, with findings based primarily on attention being directed to either ear rather than to sound content. In this experiment, a mixture of melody streams was presented to both ears identically (*diotically*) as we examined if selective auditory attention to sound content influences the ASSR. Using magnetoencephalography (MEG), we assessed the stream-specific ASSRs from three frequency-tagged melody streams when attention was directed between each melody stream, based on their respective pitch and timing. Our main results showed that selective attention enhances the ASSR power of an attended melody stream by 15 % at a general sensor level. This ability to readily capture attentional changes in a stimuli-precise manner makes the ASSR a useful tool for studying selective auditory attention, especially in complex auditory environments. Furthermore, as a secondary aim, we explored the distribution of cortical ASSR sources and their respective attentional modulation. A novel finding using distributed source modelling revealed that the ASSR is modulated by attention in many areas across the cortex, with frontal regions experiencing the strongest enhancement of up to ~ 80 %. ASSRs in the temporal and parietal cortices were enhanced by approximately 20 - 25 %. For future studies, this work can serve as a template to narrow-down possible sites of ASSR attentional modulation for further investigation.

## 1. Introduction

In light of the brain’s limited capacity to process simultaneous information, the ability to attend to one out of several competing sounds is therefore essential, allowing one to extract and process the most important information amidst a complex auditory environment. This phenomenon was first coined the “Cocktail party effect (CPE)” by Cherry in 1953 and is important to functions such as speech recognition, musicianship and threat identification^1^. In music, selective auditory attention can manifest as the ability to discern a single instrument amongst an orchestral performance, or a single voice in a choir. This ability, estimated with speech-in-noise performance^2^ and robustness of neural patterns^3^, is positively correlated with the listener’s amount of musical training^2–3^, suggesting that selective attention capabilities may be improved through strategic training regimes. While the relevance of the CPE for perception and performance is well documented, the neural mechanisms underlying this phenomenon is still not completely understood. This is partially due to the difficulties in isolating the specific brain activity that stem from one out of many simultaneous auditory sources: If you selectively attend to only the soprano voice while listening to a choir performance, how do you separate brain activity representing the soprano from that representing the rest of the choir and study how that activity is influenced by selective attention? Previous magnetoencephalography (MEG) and electroencephalography (EEG) studies on selective auditory attention have shown that time-locked neuronal activity [e.g. event-related fields (ERFs) and potentials (ERPs)] from a wide range of auditory stimuli (e.g. click, tones, speech) is increased by attention^4–7^. However, such time-locked approaches are not easily compatible with the complex and dynamic characteristics of naturalistic or continuous stimuli. Importantly, it is very difficult to distinguish between auditory sources with simultaneous onset times from their event-related activities, which is often the case in a natural auditory environment. In such scenarios, another approach using the Auditory Steady-State Response (ASSR) may be useful to isolate and assess the neural activity related to each individual sound.

The ASSR can be described as an *oscillatory* evoked potential that continuously phase-locks to the intrinsic fundamental frequency of the stimulus over the time period of stimulus presentation^8,9^. The constituent discrete frequency components of the ASSR can be retrieved from recorded MEG/EEG data using power spectral density (PSD) estimation methods such as Fourier analysis. A handy way to adjust the stimulus frequency, and consequently the ASSR frequency, while retaining much of the stimulus property (e.g. pitch, timbre) is via *amplitude modulation (AM) frequency-tagging* of the sound. This is done by increasing and decreasing the amplitude of the sound envelope (i.e. volume) at a precise rate defined by the modulation frequency (f_m_). This technique can be used to disentangle the processing of sound streams presented simultaneously, since the neural activity to each stream can be distinguished by a unique f_m_ during analysis^10–11^. In humans, the ASSR is known to reach a maximum power response at frequencies close to 40 Hz^8^, hence the term *40 Hz ASSR*. Several intermodal studies have demonstrated that the cortically generated ASSR is enhanced when attention is voluntarily directed towards (as compared to directed away from) an auditory stimuli from a competing visual stimulus^12–14^. Within the auditory domain (i.e. intramodal studies) however, results remain unclear. In some cases, selective attention tasks using dichotic stimulus presentation reported an ASSR enhancement by attention while in other cases no effect of attention was found^6, 15–17^. The inconsistency in findings suggests that whether or not attention is found to affect the ASSR depends on several experimental design factors pertaining to the stimuli, task and analytical approach. Furthermore, the majority of intramodal auditory attention ASSR studies adopt a *dichotic* experimental design wherein participants shift attention between the left and right ears, and the corresponding changes in cortical ASSRs are assessed with MEG^15,18^. Therefore, selective attention in such dichotic experiments is heavily reliant on spatial separation of the auditory input (ears) rather than perceptual separation of the sound streams based on sound content, despite the latter being an essential aspect of selective listening. Also, the spatial separation approach is inherently limited to two ears and thus only two sources, making it inapplicable to studies involving complex auditory mixtures with several sources. To the best of our knowledge, no study has examined the influence of selective attention on the 40 Hz ASSR when the same auditory mixture of multiple streams is presented to both ears (i.e. diotically), and auditory stream separation must be based solely on perceptual features of the sound content (i.e. pitch/timbre/tempo). This gap in the ASSR-attention literature may point to some challenges that researchers face in designing such an experiment, for example, in finding suitable stimuli and tasks with sufficient stream separability to evoke a detectable difference in selective attention when using diotic stimuli.

In the current study, we aim to explore this uncharted approach by using a task where selective attention is directed towards diotically-presented AM frequency-tagged melody streams that are easily differentiable by their respective timing and pitch. For the frequency-tagging, we used separate modulation frequencies at f_m_ = 37, 39, 41 Hz to individually tag each of three different melody streams, with the goal of eliciting ASSRs corresponding to the three melody streams that can be clearly separated in the frequency domain during analysis.

The primary aim of this study was to assess if ASSR power is influenced when selective attention is directed towards a specific melody stream. To assess the ASSR, we measured ongoing brain activity at millisecond temporal resolution and millimetre spatial precision using MEG^19^. At the same time, MEG is also well-suited for spatially precise modelling of brain activity at an individual anatomical level. Based on the rich literature supporting the enhancement effect of selective attention on neural signals^7, 20–23^, we hypothesized that attention increases the ASSR power corresponding to the attended stream. With sufficient signal power, we expect that this attention effect may be observed already in sensor-level data.

A secondary aim of this study is to understand the structural distribution of the cortical sources that are involved in ASSR expression and their attentional modulation. Since little is known about the source distribution of the 40 Hz ASSR, apart from its presence in the auditory cortex^24–26^, we cannot precisely point to, *a priori*, where to expect the ASSR attention effect although attention-related literature does suggest the prefrontal cortex as a likely site^27–30^. As such, we will carry out source analysis using a distributed source model to identify likely ASSR source positions, and then examine the degree to which attention modulates ASSR power in each of these ASSR source regions. In this sense, our secondary aim is more exploratory in nature and we have adopted a more data-driven approach for this part of the analysis.

## 2. Materials and Methods

### 2.1 Participants

A total of 29 participants (age 18 – 49 years, mean age = 28.0, SD = 4.9; 10 female; 2 left-handed) with normal hearing volunteered to take part in the experiment. The experiment was approved by the Regional Ethics Review Board in Stockholm (Dnr: 2017/998-31/2). Both written and oral informed consent were obtained from all participants prior to the experiment. All participants received a monetary compensation of SEK 600 (~ EUR 60). One participant was excluded due to incomplete data collection, and a second participant was excluded due to less-than-chance performance in the behavioural task, resulting in a final sample size of 27 participants for all MEG analyses.

### 2.2 Experimental Task: Melody Development Tracking (MDT) task

Participants were presented with 3 melody streams of increasing pitch [i.e. carrier frequency (f_c_) range], henceforth referred to as the Bottom voice, Middle voice, and Top voice. The participants were instructed to direct attention exclusively to the Bottom voice or Top voice according to a cue before the melodies started (e.g. “Attend bottom voice!”). At a random surprise point during melody playback, the melody stopped and participants were asked to report the latest direction of pitch change for the attended melody stream by pressing one out of three buttons, representing *falling, rising* or *constant* pitch respectively (e.g. whether the last note was falling, rising or constant relative to the note preceding it. Refer to Fig. 1). In total, 28 of these responses were collected for each participant.

**Figure 1.**
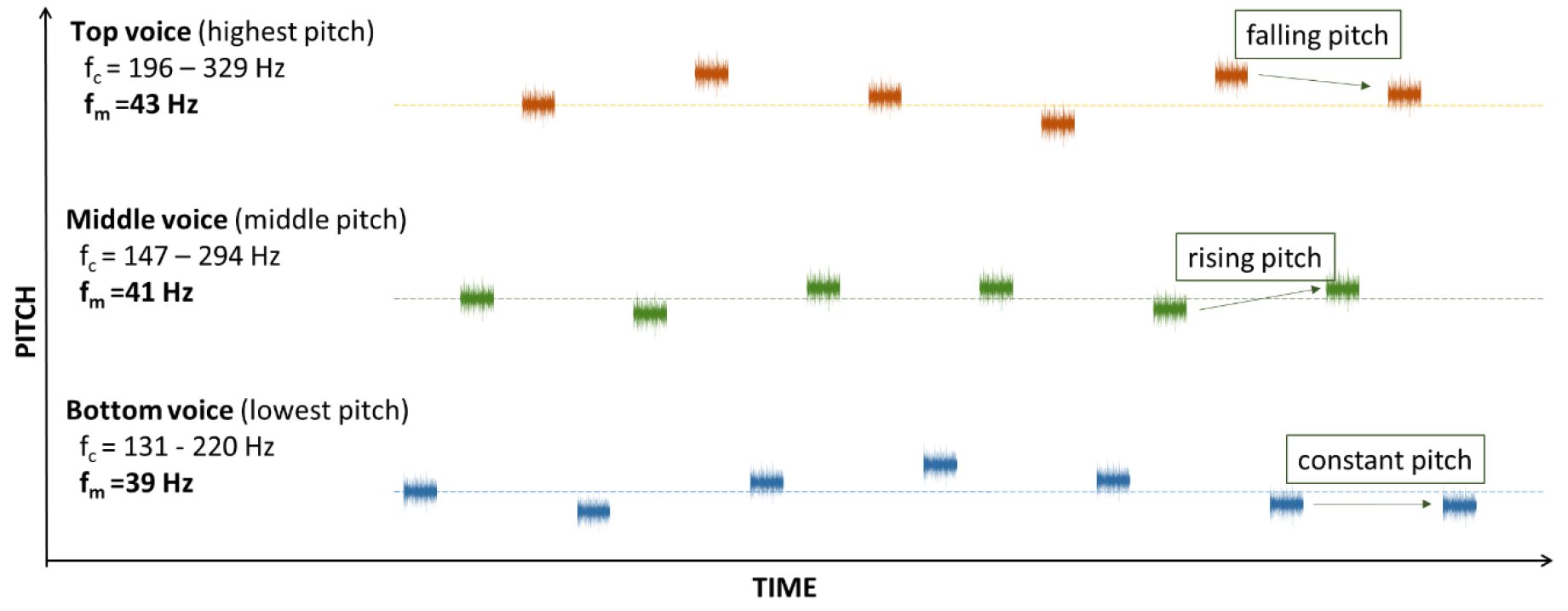
The Melody Development Tracking (MDT) task. Participants listened to three melody streams while attending to either the Bottom voice or Top voice following a cue. When the melody stopped, participants were asked to report the last direction of pitch change for the attended melody stream (i.e. falling, rising or constant pitch as illustrated). The three melody streams were presented separately in time, starting from Bottom to Top (shown in figure) or its reverse. The respective f_c_ (pitch) range and f_m_ of each stream are indicated above.

The three voices were presented separately in time, such that the voices had their note onset either in the order of Bottom-Middle-Top or its reverse, while keeping the order balanced across trials. Prior to the actual MEG recording, participants received 10 to 15 min of training to familiarize themselves with the task: Participants were deemed ready to commence with the actual experiment once they managed to report the correct answers for at least five consecutive trials. As the task was designed to require continuous selective attention to the cued melody stream, it is imperative to maintain alertness and alleviate fatigue. We therefore introduced a brief break in the task every ~5 min, during which their general attentiveness was also assessed using the Karolinska sleepiness scale^31^. To minimize movement artefacts, participants were asked not to move when listening to each melody segment, which was at most 30 s. The MEG recording time was approximately 20 min per participant, including breaks.

### 2.3 Stimuli

Each of the three voices was constructed using a stream of 750 ms long sinusoidal tones of f_c_ between 131 – 329 Hz (Bottom voice: 131 - 220 Hz; Middle voice 147 – 294 Hz; Top voice 196 – 329 Hz), generated using the Ableton Live 9 software (Berlin, Germany). At the onset and offset of each tone, we introduced a 25 ms amplitude fade-in and fade-out to avoid audible compression clicks. These tones were then amplitude-modulated sinusoidally in Ableton Live 9 using f_m_ at 39 (Bottom voice), 41 (Middle voice), and 43 (Top voice) Hz, and a modulation depth of 100% to achieve maximum ASSR power^8^. For simplicity, only tones in the C major harmonic scale were used. The duration of melody presentation was randomized to be between 9 – 30 seconds long to reduce predictability of the stop point for maintaining the participants’ attention throughout the melody. Loudness was calibrated using a soundmeter (Type 2235, Brüel & Kjær, Nærum, Denmark) to account for differences in subjective loudness for different frequency ranges^32^. The respective settings for the Bottom, Middle and Top voices were 0 dB, −6 dB and −10 dB. The stimulus was presented identically via ear tubes to both ears with the volume adjusted to be 75 dB SPL per ear, subjected to individual comfort level.

### 2.4 Data Acquisition

MEG measurements were carried out using a 306-channel whole-scalp neuromagnetometer system (Elekta TRIUX_TM_, Elekta Neuromag Oy, Helsinki, Finland). Data was recorded at a 1 kHz sampling rate, on-line bandpass filtered between 0.1-330 Hz and stored for off-line analysis. Horizontal eye-movements and eye-blinks were monitored using horizontal and vertical bipolar electroculography electrodes. Cardiac activity was monitored with bipolar electrocardiography electrodes attached below the left and right clavicle. Internal active shielding was active during MEG recordings to suppress electromagnetic artefacts from the surrounding environment. In preparation for the MEG-measurement, each participant’s head shape was digitized using a Polhemus FASTRAK. The participant’s head position and head movement were monitored during MEG recordings using head-position indicator coils. Anatomical MRIs were acquired using hi-res Sagittal T1 weighted 3D IR-SPGR (inversion recovery spoiled gradient echo) images by a GE MR750 3 Tesla scanner with the following pulse sequence parameters: 1 mm isotropic resolution, FoV 240×240 mm, acquisition matrix: 240 x 240, 180 slices 1 mm thick, bandwidth per pixel=347 Hz/pixel, Flip Angle=12 degrees, TI=400 ms, TE=2.4 ms, TR=5.5 ms resulting in a TR per slice of 1390 ms.

### 2.5 Data Processing

The acquired MEG data was pre-processed using MaxFilter (-v2.2)^33–34^,and subsequently analysed and processed using the Fieldtrip toolbox^35^ in MATLAB (Version 2016a, Mathworks Inc., Natick, MA), as well as the MNE-Python software^36^. Cortical reconstruction and volumetric segmentation of all participants’ MRI was performed with the Freesurfer image analysis suite^37^.

#### 2.5.1 Pre-Processing

MEG data was MaxFiltered by applying temporal signal space separation (tSSS) to suppress artefacts from outside the MEG helmet and to compensate for head movement during recordings^33–34^, before being transformed to a default head position. The tSSS had a buffer length of 10 s and a cut-off correlation coefficient of 0.98. The continuous MEG data was divided into 1 s-long epochs from stimulus onset (i.e. onset of each individual note). Epochs were then visually inspected for artefacts and outliers with high variance were rejected using *ft_rejectvisual*^35^. After cleaning, the remaining 69 % of all epochs were kept for further analyses. The data was divided into six experimental conditions, consisting of epochs (~100 per condition) for each of the three voices (Bottom, Middle, Top) under instructions to attend the Bottom voice or Top voice, respectively, i.e.: i) Bottom voice – Attend Bottom (Bottom-Attend), ii) Bottom voice – Attend Top (Bottom-Unattend), iii) Top voice – Attend Top (Top-Attend), iv) Top voice – Attend Bottom (Top-Unattend), v) Middle voice – Attend Bottom, vi) Middle voice – Attend Top.

#### 2.5.2 Behavioural data analysis

To assess response accuracy in the MDT task, mean task performance scores (number of correct responses out of 28 total responses) were calculated across all conditions separately for each participant.

#### 2.5.3 Sensor-space analysis

We carried out sensor-space analysis on the cleaned MEG epochs to extract the effect of selective attention on the ASSR. ERFs were also extracted to check for the manipulation of attention by the task, since it has already been well-documented in literature that attention enhances the ERF^4–7^. For these analyses, firstly, a 30 – 50 Hz bandpass filter was applied to obtain the ASSR, and a 20 Hz low-pass filter was applied to obtain the ERF. Within each participant, the filtered epochs were then averaged per condition, resulting in the *timelocked ASSR* and *timelocked ERF*. The ERF data was demeaned using an interval, 100 - 0 ms before stimulus onset, as the baseline. To acquire the ASSR power spectrum in the frequency domain, a fast Fourier transform (hanning-tapered, frequency resolution = 1 Hz) was applied to the *timelocked ASSR* data above. The ASSR power spectrum and *timelocked ERF* data were further averaged across all gradiometer sensors, after collapsing data from orthogonal planar gradiometers, to give the average gradiometer data per participant. Gradiometer sensors were selected for analysis as they are generally less noisy compared to magnetometers. The ASSR power at f_m_, (defined as 39, 41, and 43 Hz for the Bottom, Middle and Top voices respectively) was extracted accordingly for each of the six conditions to give the mean ASSR power at f_m_ per condition (e.g. For the Bottom-Attend and Bottom-Unattend conditions, the power at 39 Hz was used). To obtain the ERF sustained field amplitude per condition, the average amplitude across the *timelocked ERF* data was calculated using a 300 – 800 ms post-stimulus onset time window^6, 38^.

#### 2.5.4 Source-space analysis

In order to model the effect of selective attention on the ASSR at the anatomical level, we used a distributed source model containing 20484 dipolar sources on the cortical surface of each participant. By using a minimum-norm estimate (MNE) approach^36^, we estimated the amplitude of these sources that generated the ASSR. The *timelocked ASSR* data was used for this analysis, to produce MNE solutions for each participant that were subsequently morphed to a common head template - fsaverage. As an initial step, we calculated the group-averaged morphed MNE solution before computing its power spectral density (PSD) using Welch’s method (hanning windowed, frequency resolution = 1 Hz). We then used the middle voice (excluded from source analyses addressing the attention effect on ASSR) PSD as a localizer to identify ASSR sources across the cortex. The entire cortical sheet was divided into 105 sub-regions per hemisphere according to the Brainnetome Atlas^39^, and the PSDs of all vertices within each sub-region were averaged to give a median *localizer power* per sub-region. For the sake of clarity, we used the occipital lobe as a reference for the absence of strong and independent ASSR sources, and discarded 24 sub-regions (12 per hemisphere) with *localizer power* less than the mean *localizer power* across all vertices in the occipital lobe (4.83·10^-27^ A^2^m^2^). For each of the remaining 93 sub-regions (symmetrical across both hemispheres), PSDs of the constituent vertices were averaged to give a median PSD per Sub-region x Voice (Bottom and Top voices only) x Attend condition. Next, the power at f_m_ (i.e. the ASSR power) during Attend and Unattend conditions was extracted separately for the Bottom and Top voices. The Attend versus Unattend ASSR power difference (Attend – Unattend) for each voice was computed as a percentage of the power at the Unattend condition (*% AU change*), representing a measure of the ASSR power enhancement due to selective attention. To obtain a visual estimation of the ASSR attentional enhancement across the cortical space, we mapped the *% AU change* over all sub-regions as shown in Figure 5. For a more concise numerical representation of the attentional contrast across the brain, the 93 sub-regions were subsequently categorized into 20 regions of interests (ROIs) per hemisphere according to the Brainnetome Atlas^39^ (labels in Fig. 5). As before, the PSDs of all vertices within each ROI were median-averaged before extracting the power at f_m_ per Voice x Attend condition. The *% AU change* was computed and tabulated in Table 1, alongside the median *localizer power* per ROI.

#### 2.5.5 Point spread function of a stimulated 41 Hz ASSR at the primary auditory cortex

We used the point spread function to characterize the spatial accuracy of the source localization method by illustrating the amount of signal spread occurring due to the source estimation of a single activated source, or in this case an area of activated sources. We computed the point spread function by first simulating a continuous 41 Hz sine wave at each vertex in the primary auditory cortex (see Figure 4B inset) for both hemispheres, followed by projecting the activity to the 306 MEG sensors, and finally projecting the estimated magnetic fields back to source space using the inverse solution that was obtained in the step above. This process was repeated for each participant using their individual noise covariance matrices and head geometries, then morphed to the fsaverage head template, and eventually averaged across all 27 participants to give the final solution illustrated in Figure 4B. The power of the simulated activity was fixed at 24.1·10^-26^ A^2^m^2^, the group-averaged maximum power obtained in the MNE solution for the Middle voice localizer.

## 3. Results

### 3.1 Behavioural results

Results from the MDT task showed that overallparticipants performed significantly above the chance level of 33% (M = 67 %, SD = 21.7 %; *t*(29) = 8.30, p_one-tailed_ < 0.001). As described above, under 2.1 Participants, two participants were excluded from further analyses to give a final sample size of 27 participants for subsequent MEG analyses. MDT task performance was not significantly different between directing attention to Bottom and Top voice (p_two-tailed_ =0.92).

### 3.2 MEG results

#### 3.2.1 Sensor space

We used sensor space analysis of MEG data to evaluate our primary hypothesis: Selective attention to frequency-tagged melody streams enhances the magnitude of the ASSR corresponding to the attended stream. To extract the effect of selective attention on the ASSR for each participant, we computed the average ASSR power spectrum across gradiometer sensors for all six conditions: Bottom-Attend, Bottom-Unattend, Top-Attend, Top-Unattend, Middle voice - Attend Bottom, Middle voice – Attend Top. For each of these conditions, we also calculated the average ERF sustained field to validate that our task successfully manipulated selective attention. Figure 2 shows the across subject grand average ASSR power spectra. The ASSR peaks for each voice can be observed clearly at the respective modulation frequencies of 39 (Bottom), 41 (Middle) and 43 (Top) Hz.

**Figure 2.**
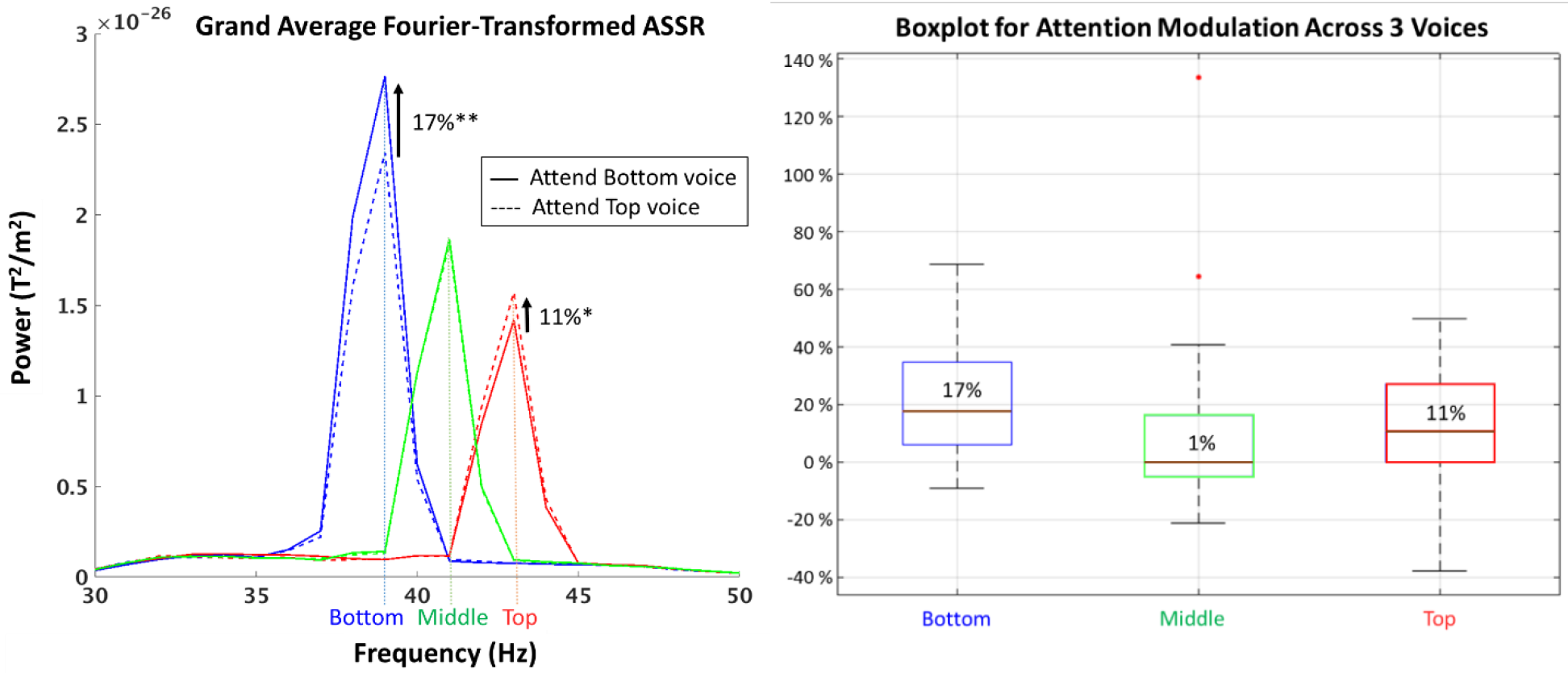
(<u>Left panel</u>) Across subject Grand Average ASSR power spectra for all conditions. ASSR power increased significantly when participants attended the corresponding Bottom (39 Hz - blue) or Top (43 Hz - red) voice. For the reference Middle voice (41 Hz - green), there was no significant difference between Attend Bottom and Attend Top. Arrows indicate median percentage attentional enhancement across all 27 participants. p<0.01**, p<0.05* (<u>Right panel</u>) Boxplot showing the distribution of all 27 participant’s percentage attentional change for the 3 voices. Median values are marked with brown lines and displayed in each box, while the bottom and top edges of each box indicated the 25 % and 75 % percentiles respectively. Outliers beyond the whiskers are plotted with red dots.

##### 3.2.1.1 Attention and ASSR power

The Attend versus Unattend contrasts, using mean power (all units are in T^2^/m^2^) at f_m_ for the Bottom and Top voices, yield significant differences with a higher power for the Attend (M_bottom_ = 2.80·10^-26^, SD_bottom_ = 3.3·10^-26^; M_top_ = 1.60·10^-26^, SD_top_ = 1.6·10^-26^) compared to Unattend (M_bottom_ = 2.39·10^-26^, SD_bottom_ = 2.7·10^-26^; M_top_ = 1.45·10^-26^, SD_top_ = 1.5·10^-26^) condition (*t*(27)_bottom_ = 3.51, p_two-tailed,bottom_ = 0.0016; *t*(27)_top_ = 2.62, p_two-tailed,top_ = 0.014). These differences are expressed as a percentage of increase relative to the Unattend condition, and indicated with arrows in Figure 2 (left panel), alongside the spread of the data across individual participants (see Fig. 2, right panel). These results confirmed our primary hypothesis that selective attention enhances ASSR power, and at an average of 14 % across both Bottom and Top voices. There was no significant difference in ASSR enhancement between the Bottom and Top voice (*t*(27) =1.22, p_two-tailed_ = 0.24). As expected, the ASSR enhancement was specific for the selectively attended voice, and was not observed on the Middle voice which participants were never instructed to attend to. Accordingly, there was no significant difference (*t*(27) =0.33, p_two-tailed_ = 0.74) between Attend Bottom (M = 1.90·10^-26^, SD = 2.0·10^-26^) and Attend Top (M = 1.89·10^-26^, SD = 2.1·10^-26^) for the Middle voice.

##### 3.2.1.2 Attention and ERFs

To validate that the MDT task manipulated attention successfully, we calculated the average ERF sustained field amplitude per Voice × Attend condition. The results from contrasting the Attend versus Unattend ERF showed significant differences for the Bottom (*t*(27) = 5.55, p_two-tailed_ < 0.001) and Top (*t*(27) = 6.27, p_two-tailed_ < 0.001) voices. As with the ASSR, for the non-attended Middle voice, there was no significant difference between Attend Bottom and Attend Top (*t*(27) = 1.18, p_two-tailed_ = 0.25). These ERF results serve as supporting evidence to show that selective attention was successfully manipulated as intended in this experiment (i.e directed exclusively to the instructed voice). The subject grand averaged ERFs per condition are illustrated in Figure 3 with arrows indicating the attentional enhancement [29 % (Bot); 26 % (Top)].

**Figure 3.**
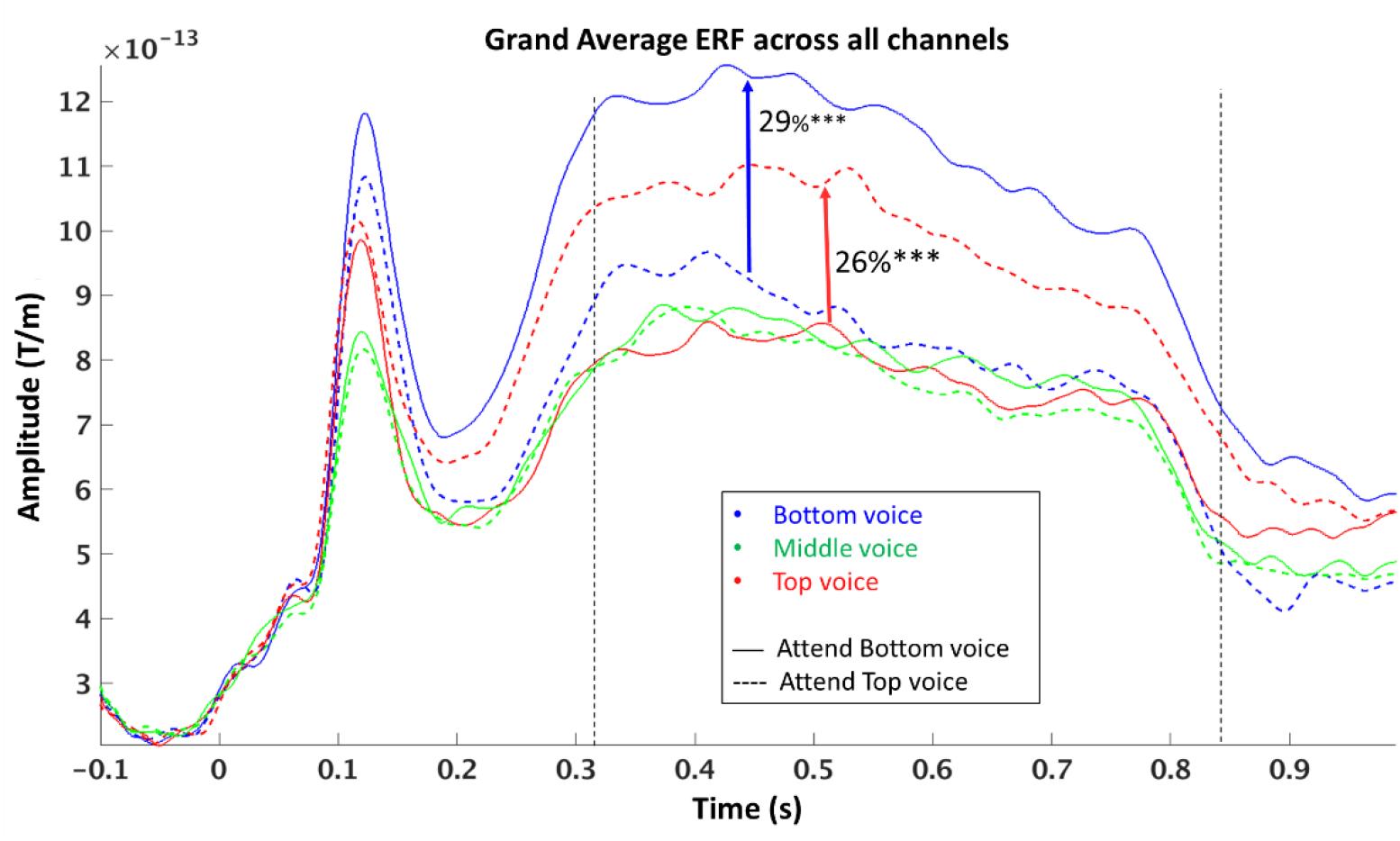
Across subject Grand Average ERF for all conditions. The amplitude of the ERF sustained field was averaged across 300-800ms post-stimulus (black vertical dashed lines) and used for comparison between Attend versus Unattend conditions. As with the ASSR results, when participants attended the Bottom (blue) or Top (red) voice, corresponding ERF amplitudes increased significantly. There was no significant difference between Middle voice – Attend Bottom and Middle voice - Attend Top (green). Arrows indicate median percentage attentional enhancement across all 27 participants. p<0.001***

#### 3.2.2 Source space

Our secondary aim to determine the cortical distribution of neural sources that are involved in ASSR expression (section 3.2.2.1 below) and their sensitivity to attentional modulation (section 3.2.2.2 below) was addressed with source space MEG analysis.

##### 3.2.2.1 Location of ASSR Sources

To identify the cortical areas involved in ASSR expression, a distributed MNE source estimate of the Middle voice localizer power was computed, revealing multiple ASSR sources that originate mainly from the temporal, parietal and frontal cortices (Figure 4A). These source positions are coherent with the results of previous studies supporting ASSR activation sites extending beyond the auditory cortex^40–42^. Unsurprisingly, sources with the strongest power were found in the primary auditory cortical regions, followed by parietal and frontal sources. In addition, the point spread function (see 2.5.5 above for details) for a simulated 41 Hz sine wave at the left and right auditory cortices was computed to assess whether the obtained ASSR sources were independent from one another or a result from a single signal in the primary auditory cortex spreading (Figure 4B). Notably, the modelled point spread function displayed a much lower maximum power at 13.7·10^-26^ A^2^m^2^, compared to the MNE solution with its maximum power at 24.1 ·10^-26^ A^2^m^2^, as well as less extensive coverage of the cortex. This shows that there exist large and systematic differences between the point spread model and our observed ASSR sources, and that the MNE source estimate cannot be explained solely by the signal spread of the primary auditory source.

**Figure 4A.**
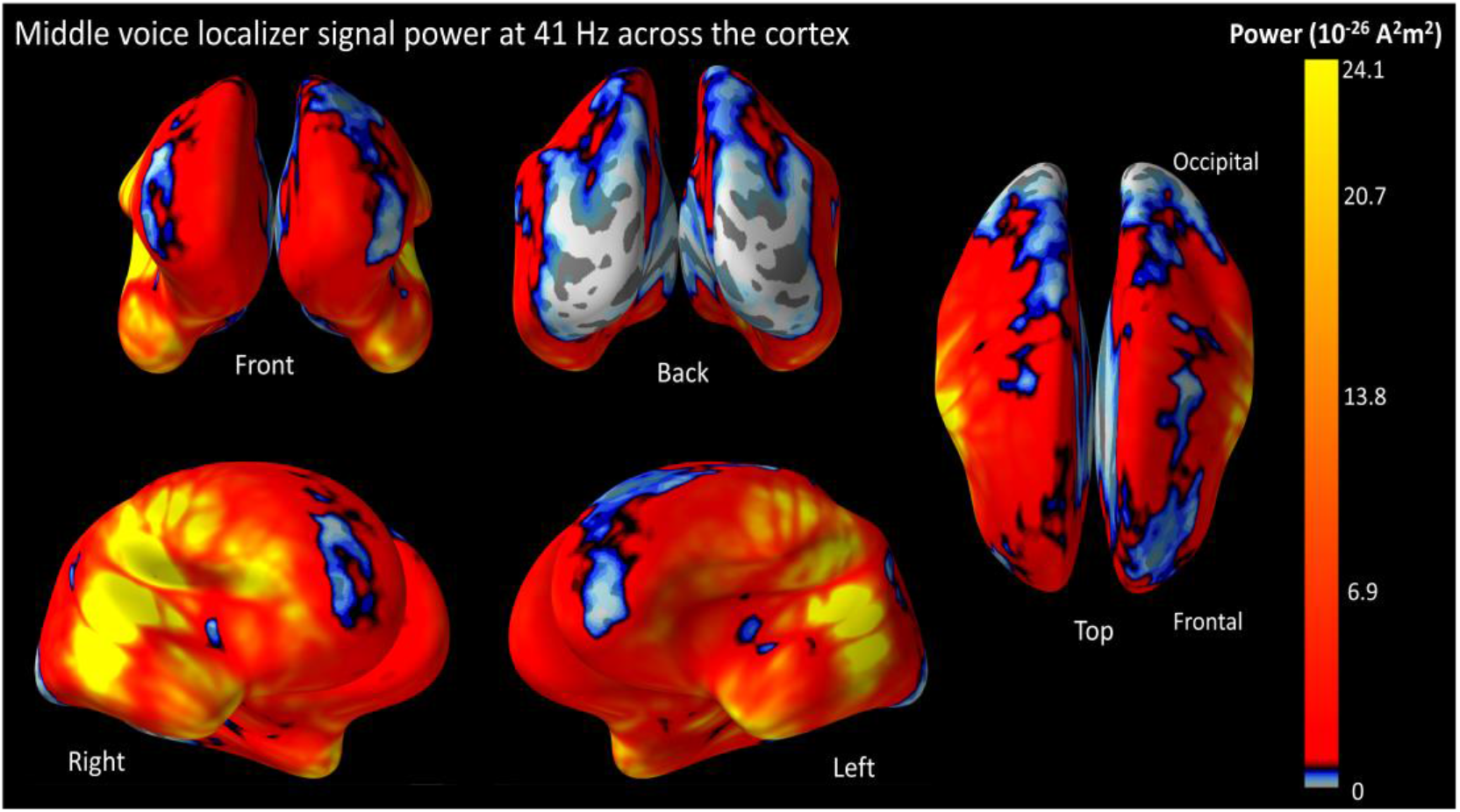
ASSR power at 41 Hz for the Middle voice localizer across the cortex. The MNE solution for the Middle voice was used to estimate the location and strength of ASSR sources. Multiple ASSR sources were found over the entire cortical sheet with the strongest located in the primary auditory cortex. Other relatively strong sources were distributed over the temporal as well as parietal cortices, while sources with moderate activity were observed in the frontal region. Overall, the ASSR was stronger in the right than left hemisphere. The strength of the ASSR is described by the colour bar on the rightmost end.

**Figure 4B.**
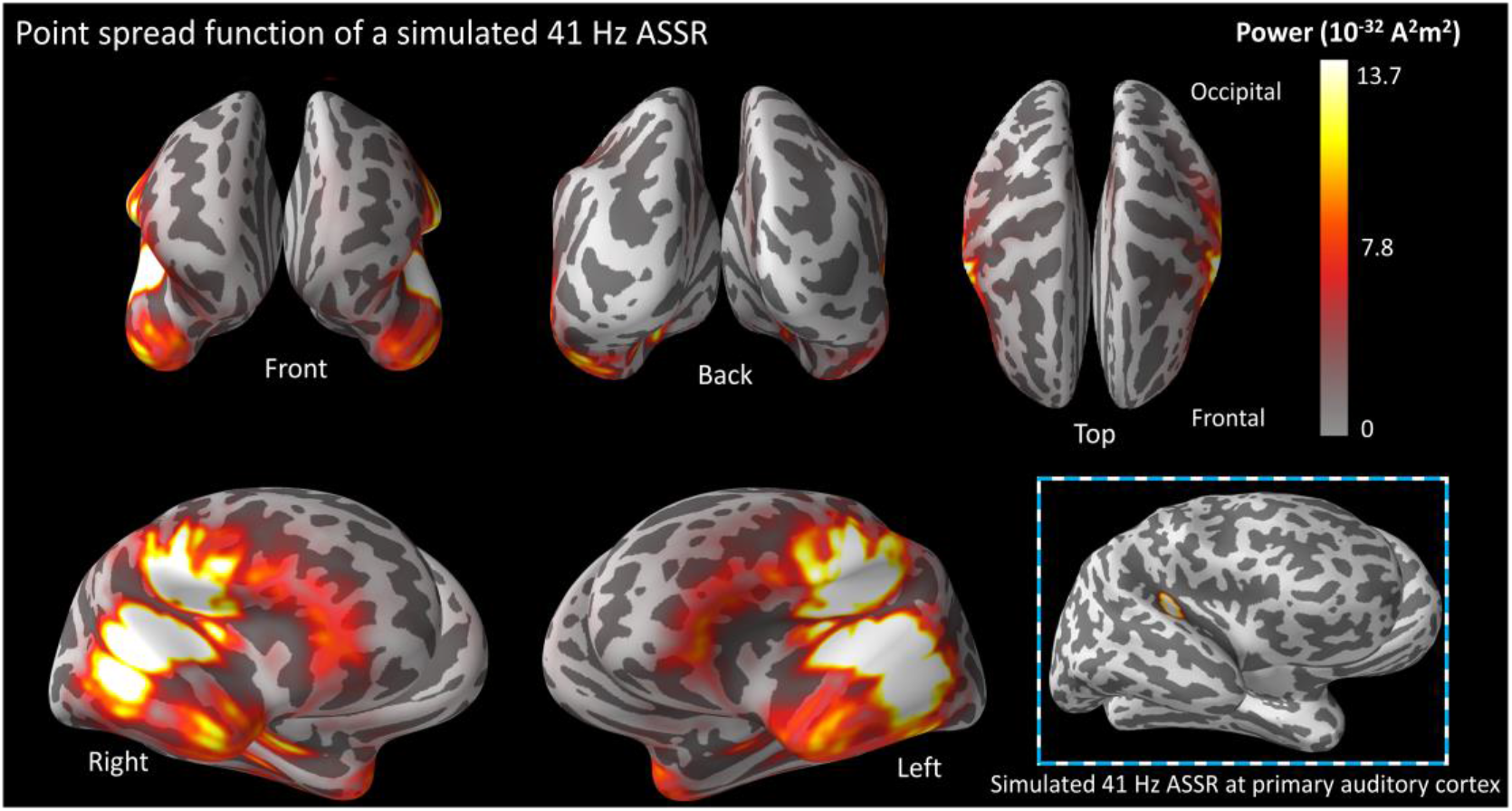
Point spread function of a simulated 41 Hz ASSR at the primary auditory cortex (both hemispheres). The power of the simulated ASSR sources (bottom-right inset) was adjusted to 24.1·10^-26^ A^2^m^2^, the maximum power of the group-averaged Middle voice localizer. After applying the inverse solution, the resultant point spread function gave a lower group-averaged maximum power of 13.7·10^-26^ A^2^m^2^.

##### 3.2.2.2 Location of ASSR Attentional Enhancement

To evaluate how much each area involved in ASSR expression is modulated by selective attention, we computed the *% AU change* - a measure of the relative ASSR attentional enhancement - across 93 sub-regions per cortical hemisphere for the Bottom and Top voices. Figure 5 shows the voice-averaged *% AU change* across these sub-regions. The frontal cortex shows a wider range of attentional modulation effects, with some focal parts exhibiting very strong attentional ASSR enhancement above 80 % (yellow) while other areas display moderately strong attentional effect around 40 % (orange). In contrast, temporal and parietal regions display weaker but more homogeneous distribution of attentional modulation across sub-regions, with ASSR enhancements typically around 20 - 25 % (dark orange).

**Figure 5.**
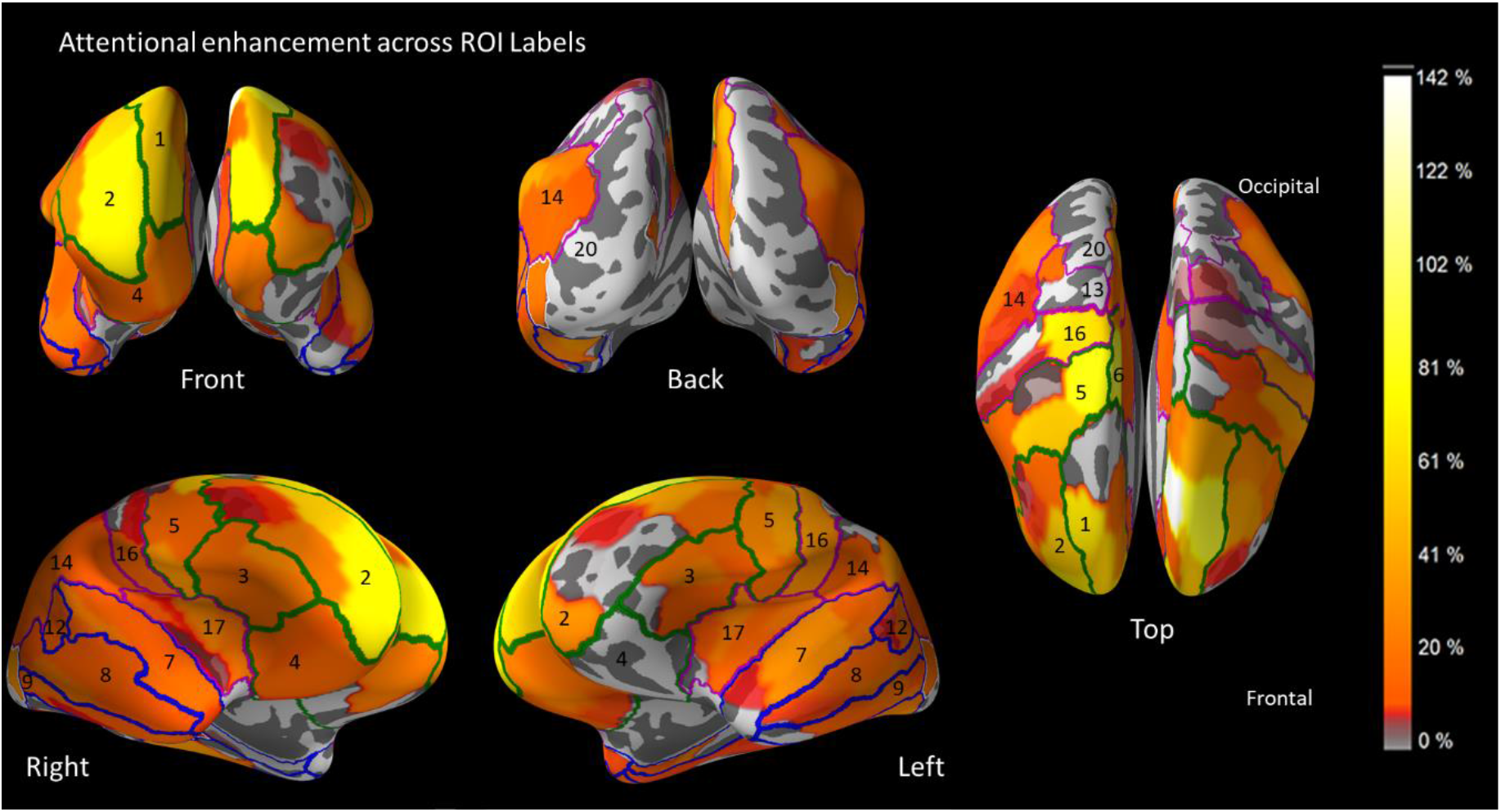
Distribution of ASSR attentional enhancement across the cortex. The average percentage increase in ASSR power between Attend and Unattend conditions across the Bottom and Top voices was computed and scaled according to the colour bar on the right. Generally speaking, frontal regions display a 2 – 4 times larger attentional enhancement than temporal and parietal regions. The frontal cortex also shows a wider range of attentional modulation effects across sub-regions, with some focal parts exhibiting above 80 % attentional ASSR enhancement (yellow) while other areas display comparatively weaker attentional effect of around 40 % (orange). On the other hand, temporal and parietal regions show more homogeneity in the distribution of attentional enhancement that revolves around 20 - 25 % (dark orange). The 93 sub-regions can be categorized into 20 ROI labels per hemisphere as demarcated above. ROIs in the frontal (green), temporal (blue), parietal (magenta) and occipital (white) lobes are numbered according to the Label # column in Table 1. Labels 10, 11, 15, 18 and 19 are located in the medial region between both hemispheres and thus not visible in this figure.

Subsequently, we categorized the sub-regions into 20 ROIs per hemisphere and compiled the *% AU change* for each in Table 1, sorted in order of decreasing median localizer power across both hemispheres (last column). The attention effect was distributed across all ROIs at an average of ~ 15 %. ROIs in the frontal gyrus appear to be most strongly and consistently enhanced by attention, with the left superior frontal gyrus (Label #1 in Tab. 1 and Fig. 5) showing up to 54 % attentional enhancement. Regions in the temporal and parietal lobes displayed up to 27 % and 35 % attentional enhancement respectively. It is useful to note that while some ROIs in the bottom rows of Table 1 have high *% AU change* (e.g. Lateral Occipital Cortex) that may suggest strong attentional enhancement, localizer ASSR power in these areas were extremely weak (within the lowest 5 % of all sub-regions for the Lateral Occipital Cortex). This calls for caution when interpreting whether the attentional enhancement in these regions stems from the presence of true ASSR sources, or is likely a spurious result from noise or field spread.

**Table 1.** % AU change for 20 ROIs, sorted in order of decreasing bi-hemispheric localizer power (rightmost column). The localizer power and % AU change are shown for the Left Hemisphere (LH), Right Hemisphere (RH) and Both Hemisphere average (BH). The first column names the lobe in which the ROI belongs, while the Label # column indicates its position numbered in Figure 5 above. Coloured rows highlight ROIs belonging to the temporal (blue), frontal (orange), parietal (pink) and occipital (white) lobes.

| Lobe | Label # | Regions of interest | % AU Change | | | Localizer Power ($A^2m^2$) | | |
| --- | --- | --- | --- | --- | --- | --- | --- | --- |
|  |  |  | LH | RH | BH | LH | RH | BH |
| Temporal | 7 | Superior temporal gyrus | 19% | 15% | 17% | 1.24E-25 | 2.25E-25 | 1.74E-25 |
| Temporal | 8 | Middle temporal gyrus | 18% | 11% | 15% | 1.14E-25 | 1.43E-25 | 1.29E-25 |
| Temporal | 12 | Posterior superior temporal sulcus | 13% | 14% | 14% | 1.06E-25 | 1.48E-25 | 1.27E-25 |
| Frontal | 3 | Inferior frontal gyrus | 15% | 18% | 16% | 6.42E-26 | 1.03E-25 | 8.36E-26 |
| Temporal | 9 | Inferior temporal gyrus | 13% | 14% | 14% | 5.87E-26 | 4.79E-26 | 5.33E-26 |
| Parietal | 16 | Postcentral gyrus | 17% | 11% | 14% | 3.84E-26 | 5.74E-26 | 4.79E-26 |
| Parietal | 17 | Insular gyrus | 14% | 15% | 15% | 2.23E-26 | 5.32E-26 | 3.77E-26 |
| Frontal | 5 | Precentral gyrus | 15% | 25% | 20% | 2.79E-26 | 4.21E-26 | 3.50E-26 |
| Temporal | 10 | Fusiform gyrus | 27% | 9% | 18% | 2.72E-26 | 3.46E-26 | 3.09E-26 |
| Temporal | 11 | Parahippocampal gyrus | -14% | 10% | -2% | 2.74E-26 | 3.17E-26 | 2.96E-26 |
| Frontal | 4 | Orbital gyrus | -8% | 15% | 3% | 2.53E-26 | 3.21E-26 | 2.87E-26 |
| Parietal | 14 | Inferior parietal lobule | 13% | 14% | 14% | 9.93E-27 | 2.23E-26 | 1.61E-26 |
| Frontal | 6 | Paracentral lobule | -5% | 59% | 27% | 1.52E-26 | 1.28E-26 | 1.40E-26 |
| Frontal | 2 | Middle frontal gyrus | 18% | 25% | 21% | 8.79E-27 | 1.41E-26 | 1.15E-26 |
| Parietal | 15 | Precuneus | -3% | 35% | 16% | 1.12E-26 | 9.25E-27 | 1.02E-26 |
| Frontal | 1 | Superior frontal gyrus | 54% | 32% | 43% | 6.67E-27 | 1.13E-26 | 8.96E-27 |
| Occipital | 20 | Lateral occipital cortex | 20% | 36% | 28% | 8.01E-27 | 8.35E-27 | 8.18E-27 |
| Parietal | 13 | Superior parietal lobule | -6% | 0% | -3% | 6.74E-27 | 9.49E-27 | 8.12E-27 |
| Occipital | 19 | MedioVentral occipital cortex | 9% | 20% | 14% | 6.23E-27 | 6.74E-27 | 6.4448E-27 |
| Parietal | 18 | Cingulate gyrus | 12% | 14% | 13% | 5.12E-27 | 7.28E-27 | 6.20E-27 |

## 4. Discussion

This study was conducted with the primary aim of examining whether selective attention to frequency-tagged melody streams (in this study coined *voices*) that are presented diotically enhances the magnitude of the ASSR specifically to the selectively attended voice. Consistent with our primary hypothesis, we observed significant enhancement of ASSR power due to selective attention in MEG sensor space. As a secondary aim, we also examined the cortical distribution of neural sources that are involved in ASSR expression and their sensitivity to attentional modulation. To this aim, we analysed the MEG data using an MNE distributed source model, and found differences in the degree of attentional enhancement across frontal, temporal and parietal ROIs, as well as between the hemispheres. While some previous studies have reported ASSR modulation when shifting selective attention between sensory modalities^12–14^ and between ears (as in dichotic listening experiments)^6, 15–17^, our study investigates this effect on diotically presented sound streams that can only be distinguished by their perceptual content (i.e. pitch and timing). This is important as content-based separation is an important part of selective auditory attention in central to functions such as speech recognition and music listening. The following section discusses the key findings and relevance of the current study.

### 4.1 Attentional enhancement of ASSR

Overall, our results showed that selective attention enhanced the 40 Hz ASSR power by an average of 15 %. We also demonstrated that this enhancement was specific to the attended Bottom and Top voices, but did not spread to the adjacent non-attended Middle voice. To the best of our knowledge, this is the first time any study has reported clear findings of ASSR attentional enhancement based solely on perceptual separation of stimuli sound content. While our results revealed stronger attentional modulation for the Bottom voice ASSR than the Top voice ASSR, we also noted that the mean Bottom voice ASSR power was higher than that of the Top voice, regardless of attentional condition. We believe that the main reason behind a lower Top voice ASSR power is that its volume was reduced to −10 dB relative to the Bottom Voice (as described under Methods). The loudness of the voices was adjusted to be subjectively equal for the MDT task, in order to compensate for the subjective amplification of higher pitch sounds in human hearing^32^, and this have created general ASSR power differences between the voices^8^. This volume difference as well as other differences between the voices, such as that in carrier frequency and modulation frequency, might also have contributed to the observed attentional differences across the Bottom and Top voices, although further studies are required to better investigate this. The modulation in ASSR power due to selective attention supports the notion of a top-down regulated gain control mechanism of attention, proposed by many authors in the past^7, 20–23^. Importantly, the results provide the first clear evidence that selective attention enhances the neuronal representation of an attended sound stream, even when the attended stream is not spatially separated from other sounds, as in dichotic listening designs.

### 4.2 Location of the ASSR and its Attentional Enhancement

Regarding the cortical distribution of ASSR sources and their sensitivity to attentional modulation, MNE results revealed sources originating from a variety of frontal, temporal and parietal regions (Fig. 4A). The accompanying point spread function (Fig. 4B) of the primary auditory cortex can be used to gauge whether these sources are truly active and independent or caused by field spread from strong sources within the primary auditory cortex. Taken together, the systematic differences between the point spread model and the observed ASSR source model favours an interpretation where frontal sources are independently active from the primary auditory sources. Although several activated regions in the parietal and secondary auditory cortices overlap with the point spread function, the much larger power generated by the MNE solution, almost twice relative to the point spread function, indicates that additional sources outside the primary auditory cortex must be present to contribute to the additional signal power. As further support for this interpretation, previous EEG^42^ and positron emission tomography (PET)^40–41^ studies have also found multiple sources generating the 40 Hz ASSR, including many regions outside the auditory pathway. These regions, especially the frontal areas, are commonly overlooked in ASSR-attention studies, which typically place exclusive focus on stronger sources within the primary auditory cortex, especially when the ASSR source is modelled as dipoles. However, as a rule of caution, we recommend readers to consider the overall distribution of ASSR source activity (Fig. 4A/B) when evaluating whether an area directly expresses an ASSR and an associated attentional modulation, or whether the observed enhancement is an indirect artefact of field spread from nearby strong sources. For example, no obvious independent ASSR sources were found in the Middle Temporal Gyrus (Label #8) and Inferior Temporal Gyrus (Label #9), leading us to believe that the observed ASSR and attentional enhancement at these areas are likely due to field spread from adjacent regions. Conversely, judging from Figure 4, the Superior Temporal Gyrus (Label #7) and Postcentral Gyrus (Label #16) both contain strongly activated and visibly independent ASSR sources, thus providing more convincing evidence that substantiates the presence of actual ASSR enhancement.

A striking finding in our source level results is that there are large differences in the degree of attentional modulation across anatomical regions, with high levels of modulation outside the auditory system. Indeed, we found that the ASSR localized to the frontal gyrus displayed the largest degree of attentional modulation. As seen in Figure 5, most cortical areas display a ~ 25 % attentional enhancement from selective attention, whereas regions in the prefrontal cortex showed up to 60 - 80 % enhancement, with the effect concentrated locally in the superior frontal gyrus. This is not surprising per se as the prefrontal cortex has been long regarded as the centre of attentional control in neuroscience literature involving auditory attention^29–30, 50–51^ as well as attention in other sensory modalities^27–28^. In addition to the frontal cortices, we also found relatively more homogeneous attentional enhancement in the temporal and parietal sub-regions of ~ 25 %. Similar to our findings, attentional enhancement of the ASSR in the auditory cortex has been reported by several studies, although limited to spatial^6, 15–17^ and intermodal^12–14^ attention. Evidence of auditory attentional modulation in the parietal cortex has also been reported in previous studies^29, 52–55^, although not within the ASSR domain, owing perhaps to the lack of documentation on ASSR sources outside the auditory cortex. Interestingly, the motor cortex, housed by the parts of the frontal and parietal lobes, is known to exhibit a robust entrainment to sensory stimulation rhythms that is also enhanced from attention^53, 56–58^. Since the ASSR may be conceptualized as an entrainment (to the stimulus) itself, it is reasonable that ASSR activity and its attentional modulation was found in the motor cortex.

### 4.3 Overcoming challenges in ASSR attentional modulation research

Since the current literature is inconsistent about whether and how intramodal auditory selective attention modulates the ASSR, a consensus on this topic has yet not been reached. This is likely attributed to factors related to stimuli, task and analytical differences. For instance, *first*, using competing stimuli with too similar properties can lead to weak perceptual separation and subsequently less effective selective attention. In many cases, the competing stimuli have similar or even identical carrier frequencies^15–16, 46^, and simultaneous onsets^47^, making it difficult for participants to differentiate between stimuli, thereby translating into a smaller ASSR power difference between Attend and Unattend conditions which the measurement instrument and analysis approach may not be sensitive enough to pick up. *Second*, several studies adopted a target detection task, placing salient targets, such as a change in frequency or intensity, in both the attended stream and distractor streams^16–17, 47^. This can result in a bottom-up effect from the distractor during the appearance of targets, thereby reducing the degree of selective attention to the attended stream. Moreover, there is evidence demonstrating that salient events amplify the ASSR in the unattended stream^48^, which can also reduce the Attend vs Unattend ASSR contrast. A *third* reason could be the narrow focus on temporal auditory core regions in source models used to localize the ASSR by most studies^15–16, 47^. Although the ASSR is strongest at these areas, a one-sided focus on these regions risks overlooking other areas such as the frontal and parietal cortices that can exhibit greater selective attention effects, as is indeed seen in our current study. In this study, we sought to alleviate these potential pitfalls by improving stream separability with the use of tones that are easily separable by timing as well as pitch, inspecting the corresponding ERFs to check for successful manipulation of selective attention, adopting a melody tracking task in place of target detection, and using a distributed source model to examine the entire cortical sheet for ASSR activity.

### 4.4 Limitations of current study

While we believe that our present results make novel contributions to the existing literature on ASSR methodology as well as to the neuroscientific understanding of selective auditory attention, the study has several limitations and calls for further work to clarify the present results. Primarily speaking, our results build on ASSR sources generated by AM frequencies close to 40 Hz and may not be generalizable across ASSRs at other frequencies as they tend to display different source distribution patterns^42^. Secondly, while the use of sine tones that are separated in time may not be an accurate representation of natural auditory mixtures such as a large choir or a symphony orchestra, the ASSR approach developed in this study is the first of its kind and serves as a stepping stone for future studies on selective attention in more natural and complex environments. Thirdly, while the sensor-space analysis provides statistical evidence affirming the attentional modulation of the ASSR (primary aim), the same cannot be said for the source-space analysis which is lacking in statistical support. Instead, the secondary aim is approached via an exploratory manner, mainly due to the lack of a-priori known sites on where to expect the attentional modulation. The resulting “brain map” (Fig. 5), constructed via several layers of averaging across trials and participants to conserve only ‘clean’ signals that were time-locked to the stimulus, is meant to serve as a nuanced illustration of ASSR attentional modulation and a guideline for future studies to demarcate possible regions of interest a-priori for statistical investigation.

### 4.5 Conclusions

In this study, we demonstrated that selective attention strongly enhances the ASSR, and that this effect can be robustly observed at sensor level. At source level, the attention effect is widely observable across the cortex and strongest in the frontal regions, which is well-aligned with current literature marking the pre-frontal cortex as the centre for attentional control^27–28, 30^. This also highlights the importance of including non-auditory areas in ASSR application studies. Overall, the current study presents clear evidence that selective auditory attention to the sound content of musical streams increases the ASSR power of the attended stream according to a specific neural pattern. Since the ASSR can readily capture these attentional changes in a stimuli-precise manner, it can serve as a useful tool for future research on selective attention in complex auditory scenarios.

## Acknowledgments

Data for this study was collected at NatMEG, the National Facility for Magnetoencephalography (http://natmeg.se), Karolinska Institutet, Sweden. The NatMEG facility is supported by Knut & Alice Wallenberg (KAW2011.0207). This study was supported by the Swedish Foundation for Strategic Research (SBE 13-0115).

## Supplementary Information

### S1 Hemispheric lateralization of the ASSR and its attentional modulation

We inspected the hemispheric lateralization of the ASSR and its attentional modulation at sensor level using the ASSR power spectrum (obtained in *Methods 2.5.3* before averaging across all sensors). The ASSR power spectrum was averaged across all gradiometer sensors within each hemisphere, to give the average ASSR power spectrum per hemisphere for each participant.

The ASSR power (all units are in T^2^/m^2^) at f_m_ = 41 Hz corresponding to the Middle voice, irrespective of Attend condition, was extracted for each hemisphere as an objective measure of the raw ASSR signal without the influence of selective attention. In line with previous findings^25, 43–45^, the ASSR power difference between Right (M = 4.44·10^-26^, SD_bottom_ = 4.2·10^-26^) and Left hemisphere (M = 2.36·10^-26^, SD_bottom_ = 2.7·10^-26^;) yield significant differences (*t*(27) = 3.76, p_two-tailed_ < 0.001), demonstrating a Right-hemispheric lateralization of the ASSR.

To check the lateralization of the ASSR attentional modulation, for each hemisphere, we extracted the ASSR power at f_m_ for the Bottom and Top voices per Attend condition. There was no significant hemispheric difference in ASSR attentional modulation for either the voices (*t*(27) =1.15, p_two-tailed,bottom_ = 0.26; *t*(27) =1.05, p_two-tailed,top_ = 0.30).

## References

1. Cherry, E. C., Some Experiments on the Recognition of Speech, with One and with Two Ears. The Journal of the Acoustical Society of America 1953, 25 (5), 975–979.

2. Parbery-Clark, A.; Skoe, E.; Lam, C.; Kraus, N., Musician enhancement for speech-in-noise. Ear and hearing 2009, 30 (6), 653.

3. Parbery-Clark, A.; Skoe, E.; Kraus, N., Musical Experience Limits the Degradative Effects of Background Noise on the Neural Processing of Sound. The Journal of Neuroscience 2009, 29 (45), 14100–14107.

4. Hansen, J. C.; Dickstein, P. W.; Berka, C.; Hillyard, S. A., Event-related potentials during selective attention to speech sounds. Biological Psychology 1983, 16 (3–4), 211–224.

5. Woods, D. L.; Hillyard, S. A.; Hansen, J. C., Event-related brain potentials reveal similar attentional mechanisms during selective listening and shadowing. Journal of Experimental Psychology: Human Perception and Performance 1984, 10 (6), 761–777.

6. Bidet-Caulet, A.; Fischer, C.; Besle, J.; Aguera, P.-E.; Giard, M.-H.; Bertrand, O., Effects of Selective Attention on the Electrophysiological Representation of Concurrent Sounds in the Human Auditory Cortex. The Journal of Neuroscience 2007, 27 (35), 9252–9261.

7. Woldorff, M. G.; Gallen, C. C.; Hampson, S. A.; Hillyard, S. A.; Pantev, C.; Sobel, D.; Bloom, F. E., Modulation of early sensory processing in human auditory cortex during auditory selective attention. Proceedings of the National Academy of Sciences of the United States of America 1993, 90 (18), 8722–8726.

8. Ross, B.; Borgmann, C.; Draganova, R.; Roberts, L. E.; Pantev, C., A high-precision magnetoencephalographic study of human auditory steady-state responses to amplitude-modulated tones. J Acoust Soc Am 2000, 108 (2), 679–91.

9. Regan, D., Human brain electrophysiology: evoked potentials and evoked magnetic fields in science and medicine. Elsevier: New York, 1989.

10. Picton, T. W.; John, M. S.; Dimitrijevic, A.; Purcell, D., Human auditory steady-state responses. Int J Audiol 2003, 42 (4), 177–219.

11. Lins, O. G.; Picton, T. W., Auditory steady-state responses to multiple simultaneous stimuli. Electroencephalogr Clin Neurophysiol 1995, 96 (5), 420–32.

12. Gander, P. E.; Bosnyak, D. J.; Roberts, L. E., Evidence for modality-specific but not frequency-specific modulation of human primary auditory cortex by attention. Hearing Research 2010, 268 (1), 213–226.

13. Saupe, K.; Schröger, E.; Andersen, S. K.; Müller, M. M., Neural Mechanisms of Intermodal Sustained Selective Attention with Concurrently Presented Auditory and Visual Stimuli. Frontiers in Human Neuroscience 2009, 3, 58.

14. Keitel, C.; Schröger, E.; Saupe, K.; Müller, M. M., Sustained selective intermodal attention modulates processing of language-like stimuli. Experimental Brain Research 2011, 213 (2), 321–327.

15. Müller, N.; Schlee, W.; Hartmann, T.; Lorenz, I.; Weisz, N., Top-Down Modulation of the Auditory Steady-State Response in a Task-Switch Paradigm. Frontiers in Human Neuroscience 2009, 3, 1.

16. Bharadwaj, H.; Lee, A. K. C.; Shinn-Cunningham, B., Measuring auditory selective attention using frequency tagging. Frontiers in Integrative Neuroscience 2014, 8 (6).

17. Lazzouni, L.; Ross, B.; Voss, P.; Lepore, F., Neuromagnetic auditory steady-state responses to amplitude modulated sounds following dichotic or monaural presentation. Clinical Neurophysiology 2010, 121 (2), 200–207.

18. Bharadwaj, H. M.; Lee, A. K. C.; Shinn-Cunningham, B. G., Measuring auditory selective attention using frequency tagging. Frontiers in Integrative Neuroscience 2014, 8, 6.

19. Cohen, D., Magnetoencephalography: detection of the brain’s electrical activity with a superconducting magnetometer. Science 1972, 175 (4022), 664–6.

20. Hillyard, S. A.; Vogel, E. K.; Luck, S. J., Sensory gain control (amplification) as a mechanism of selective attention: electrophysiological and neuroimaging evidence. Philosophical transactions of the Royal Society of London. Series B, Biological sciences 1998, 353 (1373), 1257–70.

21. Hillyard, S. A.; Hink, R. F.; Schwent, V. L.; Picton, T. W., Electrical Signs of Selective Attention in the Human Brain. Science 1973, 182 (4108), 177–180.

22. Kerlin, J. R.; Shahin, A. J.; Miller, L. M., Attentional Gain Control of Ongoing Cortical Speech Representations in a “Cocktail Party”. The Journal of Neuroscience 2010, 30 (2), 620–628.

23. Lee, A. K. C.; Larson, E.; Maddox, R. K.; Shinn-Cunningham, B. G., Using neuroimaging to understand the cortical mechanisms of auditory selective attention. Hearing Research 2014, 307, 111–120.

24. Schoonhoven, R.; Boden, C. J. R.; Verbunt, J. P. A.; de Munck, J. C., A whole head MEG study of the amplitude-modulation-following response: phase coherence, group delay and dipole source analysis. Clinical Neurophysiology 2003, 114 (11), 2096–2106.

25. Steinmann, I.; Gutschalk, A., Potential fMRI correlates of 40-Hz phase locking in primary auditory cortex, thalamus and midbrain. NeuroImage 2011, 54 (1), 495–504.

26. Lamminmäki, S.; Parkkonen, L.; Hari, R., Human Neuromagnetic Steady-State Responses to Amplitude-Modulated Tones, Speech, and Music. Ear and Hearing 2014, 35 (4), 461–467.

27. Cohen, R. A., Attention and the Frontal Cortex. In The Neuropsychology of Attention, Springer US: Boston, MA, 2014; pp 335–379.

28. Foster, J. K.; Eskes, G. A.; Stuss, D. T., The cognitive neuropsychology of attention: A frontal lobe perspective. Cognitive Neuropsychology 1994, 11 (2), 133–147.

29. Shomstein, S.; Yantis, S., Parietal Cortex Mediates Voluntary Control of Spatial and Nonspatial Auditory Attention. The Journal of Neuroscience 2006, 26 (2), 435–439.

30. Plakke, B.; Romanski, L. M., Auditory connections and functions of prefrontal cortex. Frontiers in neuroscience 2014, 8, 199–199.

31. Akerstedt, T.; Gillberg, M., Subjective and objective sleepiness in the active individual. The International journal of neuroscience 1990, 52 (1-2), 29–37.

32. Robinson, D. W.; Dadson, R. S., A re-determination of the equal-loudness relations for pure tones. British Journal of Applied Physics 1956, 7 (5), 166.

33. Taulu, S.; Kajola, M.; Simola, J., Suppression of Interference and Artefacts by the Signal Space Separation Method. 2004; Vol. 16, p 269–75.

34. Taulu, S.; Simola, J., Spatiotemporal signal space separation method for rejecting nearby interference in MEG measurements. Physics in medicine and biology 2006, 51 (7), 1759–68.

35. Oostenveld, R.; Fries, P.; Maris, E.; Schoffelen, J.-M., FieldTrip: Open Source Software for Advanced Analysis of MEG, EEG, and Invasive Electrophysiological Data. Computational Intelligence and Neuroscience 2011, 2011, 9.

36. Gramfort, A.; Luessi, M.; Larson, E.; Engemann, D.; Strohmeier, D.; Brodbeck, C.; Goj, R.; Jas, M.; Brooks, T.; Parkkonen, L.; Hämäläinen, M., MEG and EEG data analysis with MNE-Python. Frontiers in Neuroscience 2013, 7 (267).

37. Fischl, B., FreeSurfer. Neuroimage 2012, 62 (2), 774–81.

38. Picton, T. W.; Woods, D. L.; Proulx, G. B., Human auditory sustained potentials. I. The nature of the response. Electroencephalography and clinical neurophysiology 1978, 45 (2), 186–197.

39. Fan, L.; Chu, C.; Li, H.; Chen, L.; Xie, S.; Zhang, Y.; Yang, Z.; Jiang, T.; Laird, A. R.; Wang, J.; Zhuo, J.; Yu, C.; Fox, P. T.; Eickhoff, S. B., The Human Brainnetome Atlas: A New Brain Atlas Based on Connectional Architecture. Cerebral Cortex 2016, 26 (8), 3508–3526.

40. Reyes, S. A.; Lockwood, A. H.; Salvi, R. J.; Coad, M. L.; Wack, D. S.; Burkard, R. F., Mapping the 40-Hz auditory steady-state response using current density reconstructions. Hearing Research 2005, 204 (1), 1–15.

41. Reyes, S. A.; Salvi, R. J.; Burkard, R. F.; Coad, M. L.; Wack, D. S.; Galantowicz, P. J.; Lockwood, A. H., PET imaging of the 40 Hz auditory steady state response. Hearing Research 2004, 194 (1), 73–80.

42. Farahani, E. D.; Goossens, T.; Wouters, J.; van Wieringen, A., Spatiotemporal reconstruction of auditory steady-state responses to acoustic amplitude modulations: Potential sources beyond the auditory pathway. NeuroImage 2017, 148, 240–253.

43. Ross, B.; Herdman, A. T.; Pantev, C., Right Hemispheric Laterality of Human 40 Hz Auditory Steady-state Responses. Cerebral Cortex 2005, 15 (12), 2029–2039.

44. Schneider, P.; Scherg, M.; Dosch, H. G.; Specht, H. J.; Gutschalk, A.; Rupp, A., Morphology of Heschl’s gyrus reflects enhanced activation in the auditory cortex of musicians. Nat Neurosci 2002, 5 (7), 688–694.

45. Weisz, N.; Lithari, C., Amplitude modulation rate dependent topographic organization of the auditory steady-state response in human auditory cortex. Hearing Research 2017, 354, 102–108.

46. Mahajan, Y.; Davis, C.; Kim, J., Attentional Modulation of Auditory Steady-State Responses. PLoS ONE 2014, 9 (10), e110902.

47. Gander, P. E.; Bosnyak, D. J.; Roberts, L. E., Evidence for modality-specific but not frequency-specific modulation of human primary auditory cortex by attention. Hearing Research 2010, 268 (1), 213–226.

48. Shuai, L.; Elhilali, M., Task-dependent neural representations of salient events in dynamic auditory scenes. Frontiers in Neuroscience 2014, 8 (203).

49. Xiang, J.; Simon, J.; Elhilali, M., Competing Streams at the Cocktail Party: Exploring the Mechanisms of Attention and Temporal Integration. The Journal of Neuroscience 2010, 30 (36), 12084–12093.

50. Pugh, K. R.; Shaywitz, B. A.; Shaywitz, S. E.; Fulbright, R. K.; Byrd, D.; Skudlarski, P.; Shankweiler, D. P.; Katz, L.; Constable, R. T.; Fletcher, J.; Lacadie, C.; Marchione, K.; Gore, J. C., Auditory Selective Attention: An fMRI Investigation. NeuroImage 1996, 4 (3), 159–173.

51. Tzourio, N.; El Massioui, F.; Crivello, F.; Joliot, M.; Renault, B.; Mazoyer, B., Functional Anatomy of Human Auditory Attention Studied with PET. NeuroImage 1997, 5 (1), 63–77.

52. Wu, C. T.; Weissman, D. H.; Roberts, K. C.; Woldorff, M. G., The neural circuitry underlying the executive control of auditory spatial attention. Brain Res 2007, 1134 (1), 187–98.

53. Besle, J.; Schevon, C. A.; Mehta, A. D.; Lakatos, P.; Goodman, R. R.; McKhann, G. M.; Emerson, R. G.; Schroeder, C. E., Tuning of the Human Neocortex to the Temporal Dynamics of Attended Events. The Journal of Neuroscience 2011, 31 (9), 3176–3185.

54. Alain, C.; Arnott, S. R.; Hevenor, S.; Graham, S.; Grady, C. L., “What” and “where” in the human auditory system. Proceedings of the National Academy of Sciences 2001, 98 (21), 12301–12306.

55. Hill, K. T.; Miller, L. M., Auditory Attentional Control and Selection during Cocktail Party Listening. Cerebral Cortex (New York, NY) 2010, 20 (3), 583–590.

56. Horton, C.; D’Zmura, M.; Srinivasan, R., Suppression of competing speech through entrainment of cortical oscillations. Journal of neurophysiology 2013, 109 (12), 3082–3093.

57. Fujioka, T.; Ross, B.; Trainor, L. J., Beta-Band Oscillations Represent Auditory Beat and Its Metrical Hierarchy in Perception and Imagery. The Journal of Neuroscience 2015, 35 (45), 15187–15198.

58. Sameiro-Barbosa, C. M.; Geiser, E., Sensory Entrainment Mechanisms in Auditory Perception: Neural Synchronization Cortico-Striatal Activation. Frontiers in Neuroscience 2016, 10 (361).

